# Integrating population genetics to define conservation units from the core to the edge of *Rhinolophus ferrumequinum* western range

**DOI:** 10.1101/662643

**Authors:** Orianne Tournayre, Jean-Baptiste Pons, Maxime Leuchtmann, Raphael Leblois, Sylvain Piry, Ondine Filippi-Codaccioni, Anne Loiseau, Jeanne Duhayer, Inazio Garin, Fiona Mathews, Sébastien Puechmaille, Nathalie Charbonnel, Dominique Pontier

**Affiliations:** CBGP, INRA, CIRAD, IRD, Montpellier SupAgro, Université de Montpellier, Montferrier-sur-Lez Cedex, France; LabEx ECOFECT « Ecoevolutionary Dynamics of Infectious Diseases », Université de Lyon, Lyon, France; Nature Environnement, Surgères, France; Department of Zoology and Animal Cell Biology, University of the Basque Country, Bilbao, the Basque Country, Spain; CNRS, Laboratoire de Biométrie et Biologie évolutive, UMR5558, Université de Lyon, Université Lyon 1, Villeurbanne, France; College of Life Sciences, University of Sussex, Falmer, UK; ISEM, Univ Montpellier, CNRS, EPHE, IRD, Montpellier, France; Groupe Chiroptères de Midi-Pyrénées (CREN-GCMP), Toulouse, France

**Keywords:** Chiroptera, microsatellites, population genetics, connectivity, conservation, demographic inference

## Abstract

The greater horseshoe bat *(Rhinolophus ferrumequinum)* is among the most widespread bat species in Europe but it has experienced severe declines, especially in Northern Europe. This species is listed Near Threatened in the European *IUCN Red List of Threatened Animals* and it is considered to be highly sensitive to human activities and particularly to habitat fragmentation. Therefore, understanding the population boundaries and demographic history of populations of this species is of primary importance to assess relevant conservation strategies. In this study, we used 17 microsatellite markers to assess the genetic diversity, the genetic structure and the demographic history of *R. ferrumequinum* colonies in the Western European part of its distribution. We found high levels of genetic diversity and large population size on the European mainland and lower estimates in England and Northern France. Analyses of clustering and isolation by distance showed a barrier effect of the Channel and potentially of the Mediterranean Sea on *R. ferrumequinum* bat dispersal. Conversely, we could not reveal any gene flow disruption from both sides of the Western Pyrenees. These results provide important information to improve the delineation of *R. ferrumequinum* management units in its western range. We suggest that a large management unit corresponding to the European mainland population must be considered. Particular attention should be given to mating territories as they seem to play a key role in maintaining the high levels of genetic mixing between colonies. Smaller management units corresponding to English and Northern France colonies must also be implemented. These insular or peripheral colonies could be at higher risk of extinction in a near future.

## Introduction

Biodiversity is dramatically declining at an accelerating rate for most animal groups (Butchart et al., 2010; Hoffmann et al., 2010; Sánchez-Bayo & Wyckhuys, 2019). According to the IUCN Red List, more than 26,500 species (27% of all assessed species) are threatened with extinction. This phenomenon results from a combination of ecological factors (e.g. habitat fragmentation and destruction, pollution, introduction of invasive species, climate change) that affect population’ sizes and connectivity. As a consequence, these populations become strongly exposed to the negative impacts of inbreeding and genetic drift (Frankham, 2005). Preserving the genetic diversity of such small and isolated populations is therefore essential to avoid inbreeding depression, to maintain genetic variability that may be useful for adaptation, in particular in response to environmental changes, and ultimately, to promote population persistence (Reed and Frankham, 2003). To address this issue and before drawing efficient conservation programs, an important pre-requisite is to gather knowledge on population boundaries and demography. In recent decades, population genetics has been combined with more classical ecological studies to infer population demographic features, including the detection of recent demographic declines or the quantification of connectivity between populations (e.g. Vignaud et al., 2014; Vonhof and Russell, 2015). Ultimately, these population genetics studies may enable to delineate functional and evolutionary conservation units such as ‘Management Units’, which are appropriate for species monitoring and management. This concept of ‘Management Units’ was proposed by Moritz (1994), and relies on the hypothesis that genetically differentiated populations should have a high priority for separate management and conservation. Defining appropriate management units is far from being trivial. Too large units may increase the risk of extinction of “cryptic” populations that would require specific strategies. On the contrary, splitting a large population in different conservation units with different strategies may lead to excessive management strategies beyond requirements, or to inappropriate strategies limiting connectivity for example.

The application of population genetics to such issues of conservation biology has been especially important for endangered species that are difficult to monitor with ecological methods, such as bats. Indeed, bats are very sensitive to climate change and human activities (Jones, Jacobs, Kunz, Willig, & Racey, 2009; Voigt & Kingston, 2016). Almost one quarter of bat species in the world are considered to be Threatened and another quarter as Near Threatened (Mickleburgh, Hutson, & Racey, 2002). Temperate-zone bats are nocturnal, small, highly mobile and the location of their roosts are often poorly known, characteristics that makes monitoring and assessment of their extinction risk difficult (O’Shea, Bogan, & Ellison, 2003). Conservation programs are often established considering local and national scales. Unfortunately, these scales are usually not defined on the basis of biological knowledge on population delineation and demography, but instead conform to administrative borders that rarely correspond to natural ecological boundaries. This is likely to limit the efficiency and coherence of conservation strategies. Population genetics might therefore enable to improve the definition of appropriate management units of bat populations (e.g. Dool et al., 2016a; Ibouroi et al., 2018). Besides, demographic inferences based on population genetics may be particularly relevant to highlight the need of conservation management for bat species. As such, Durrant et al. (2009) have been able to evidence a recent decline and high levels of inbreeding in British populations of Bechstein’s bat *(Myotis bechsteinii).*

Among European bat species, the greater horseshoe bat *(Rhinolophus ferrumequinum)* is particularly interesting to address conservation issues from population genetics. This insectivorous species – which seasonally uses hibernation and maternity roosts – has experienced dramatic declines, particularly in Northern Europe (e.g. Belgium, Luxembourg, England) where it is now considered rare or extinct (Battersby, 2005; Kervyn, Lamotte, Nyssen, & Verschuren, 2009; Pir, 2009). In some countries, such as the UK, there is evidence of recent population increases, but these need to be placed in the context of catastrophic historical declines (Mathews et al., 2018). The species is included in Appendix II of Bern Convention, Appendix II of the Bonn Convention, Annex II and Annex IV of the Directive on the conservation of Natural Habitat and of Wild Fauna and Flora and is listed in the *IUCN Red List of Threatened Animals* (International Union for the Conservation of Nature, 2017). The reasons for the disappearance of the populations of *R. ferrumequinum* are difficult to identify. It is therefore important to implement conservation programs, at adequate geographical scales and based on a solid knowledge of *R. ferrumequinum* population dynamics.

Previous phylogeographic studies of *R. ferrumequinum* have revealed a unique genetic cluster in Western Europe mainland, ranging from Portugal to Italy, that resulted from the expansion of a single population originating from a Western Asian refugium (Flanders et al. 2009 ? Rossiter et al., 2007). Despite a lack of genetic differentiation observed over thousands of kilometers on the continent, strong genetic differentiation was observed over tens to several hundreds of kilometers within the United-Kingdom. This result underlined the importance of sampling density in population genetics studies to detect finer genetic pattern and particular population functioning (e.g. source-sink dynamics). It is therefore particularly relevant to reinforce the sampling in France, where *R. ferrumequinum* is considered as Near Threatened. Indeed the distribution of *R. ferrumequinum* is very disparate there and previous phylogeographic studies have not considered the genetic variability and structure in this country. This patchy distribution implies potential differences in population size, connectivity levels and therefore exposition to extinction risk. More specifically, most of the known roosts of *R. ferrumequinum* in France are located on the Atlantic coast (Vincent and Bat Group SFEPM, 2014). The 4^th^ largest hibernating population (about 7000 individuals) and the 10^th^ largest summer population (about 2000 individuals) are found in the Poitou-Charentes region. This region has therefore a strong conservation responsibility to preserve this species and several biological programs have been dedicated to *R. ferrumequinum* study at this scale. *R. ferrumequinum* population dynamics is being assessed using counts of individuals performed in hibernation roosts by local naturalist associations over the three decades. These surveys have emphasised a decline of *R. ferrumequinum* population in this region (Poitou-Charentes Nature, 2017). *R. ferrumequinum* experiences important seasonal fluctuations in abundance that reflect seasonal changes of roosts enabling to fill appropriate abiotic conditions such as temperature or hygrometry. These fluctuations suggest large movements and potential gene flow of bats during winter and summer. However, we still have only limited knowledge about connectivity and genetic mixing between French *R. ferrumequinum* colonies.

In this study, we proposed to identify population boundaries of *R. ferrumequinum* in its western geographic distribution range and to infer the demographic history of these populations. More specifically, we aimed at assessing the biological relevance of considering the French Poitou-Charentes region as a singular Management Unit (MU), as it is currently done on the basis of political constraints. We combined several population genetics approaches based on a dense sampling of *R. ferrumequinum* maternity colonies in the Poitou-Charentes region and we also added further samples in France, Spanish Basque country, England and Tunisia. This sampling scheme in growing circle encompasses the northern edge of *R. ferrumequinum* distribution, and enabled us to question the potential impact of a variety of large geographic barriers (mountains and seas) on the connectivity of *R. ferrumequinum* colonies. We estimated the genetic diversity and effective size of colonies to evaluate their ‘genetic health’ and infer conservation priority. We expected lower levels of genetic diversity at the edge of *R. ferrumequinum* distribution range. In addition we examined patterns of genetic differentiation to evaluate the connectivity between *R. ferrumequinum* colonies. We expected a disruption of gene flow between colonies located either side of the Mediterranean or Channel seas, as it has already been shown that sea is a barrier to dispersal for several bat species (Castella et al., 2000; García-Mudarra, Ibáñez, & Juste, 2009). We also questioned the impact of the Pyrenees as a potential geographical barrier on *R. ferrumequinum* gene flow as this species is usually confined to lowlands (<1000m elevation) (Le Roux et al., 2017). However, we did not expect a strong disruption of gene flow due to the Pyrenees between French and Spanish Basque colonies as observed in previous studies (Rossiter et al., 2007).

Altogether, our results should provide important information about *R. ferrumequinum* population genetics that will complement ecological knowledge gathered by local naturalist associations to define appropriate management units in Western Europe. Both academic and non-academic partners were involved in this work to guarantee that the results would directly inform conservation and management action (Britt, Haworth, Johnson, Martchenko, & Shafer, 2018). More broadly, this study illustrates how population genetics may bring important information to delineate bat management units and to design conservation programs of bat species at relevant geographical scales.

## Material and methods

### Ethical statements

Authorization for bat capture in France was provided by the Ministry of Ecology, Environment, and Sustainable development over the period 2015-2020 (approval no. C692660703 from the Departmental Direction of Population Protection (DDPP, Rhone, France). All methods were approved by the Museum National d’Histoire Naturelle (MNHN) and the Société Française pour I’Étude et la Protection des Mammifères (SFEPM). Authorization for bat captures in Spanish Basque Country was provided by the corresponding regional Ministries and Councils. Capture and handling protocols followed published guidelines for treatment of animals in research and teaching (Buchanan et al., 2012) and were approved by the Ethics Committee at the University of the Basque Country (Ref. CEBA/219/2012/GARIN ATORRASAGASTI). Authorisation for capture in England was given by Natural England (Ref. 2017-29766-SCI-SCI) and sampling was undertaken under licence from the Home Office (PPL 3003431).

### Biological material

This study is based on 864 greater horseshoe bats that were captured using harp-traps set up in 24 maternity colonies in France (summer 2016 to 2018), 28 captured in one maternity colony in the Spanish Basque Country (summer 2012), 36 captured in two maternity colonies in England (summer 2018) and 22 captured in one maternity colony in Tunisia (summer 2012). Details are provided in Table 1 and Figure 1. Maternity colonies correspond to the roosts where female bats gather from May to August, give birth and rear their young (Fenton, 1983; Ransome & Hutson, 2000). This sampling leads to unbalanced sex and age ratios, with respectively 887 females, 62 males and one undetermined, 808 adults, 130 juveniles (less than two years old) and 12 undetermined. Distances between sampling colonies varied from 2.53 km (closest colonies from Western France) to 1,830 km (colonies from England and Tunisia).

**Figure 1.**
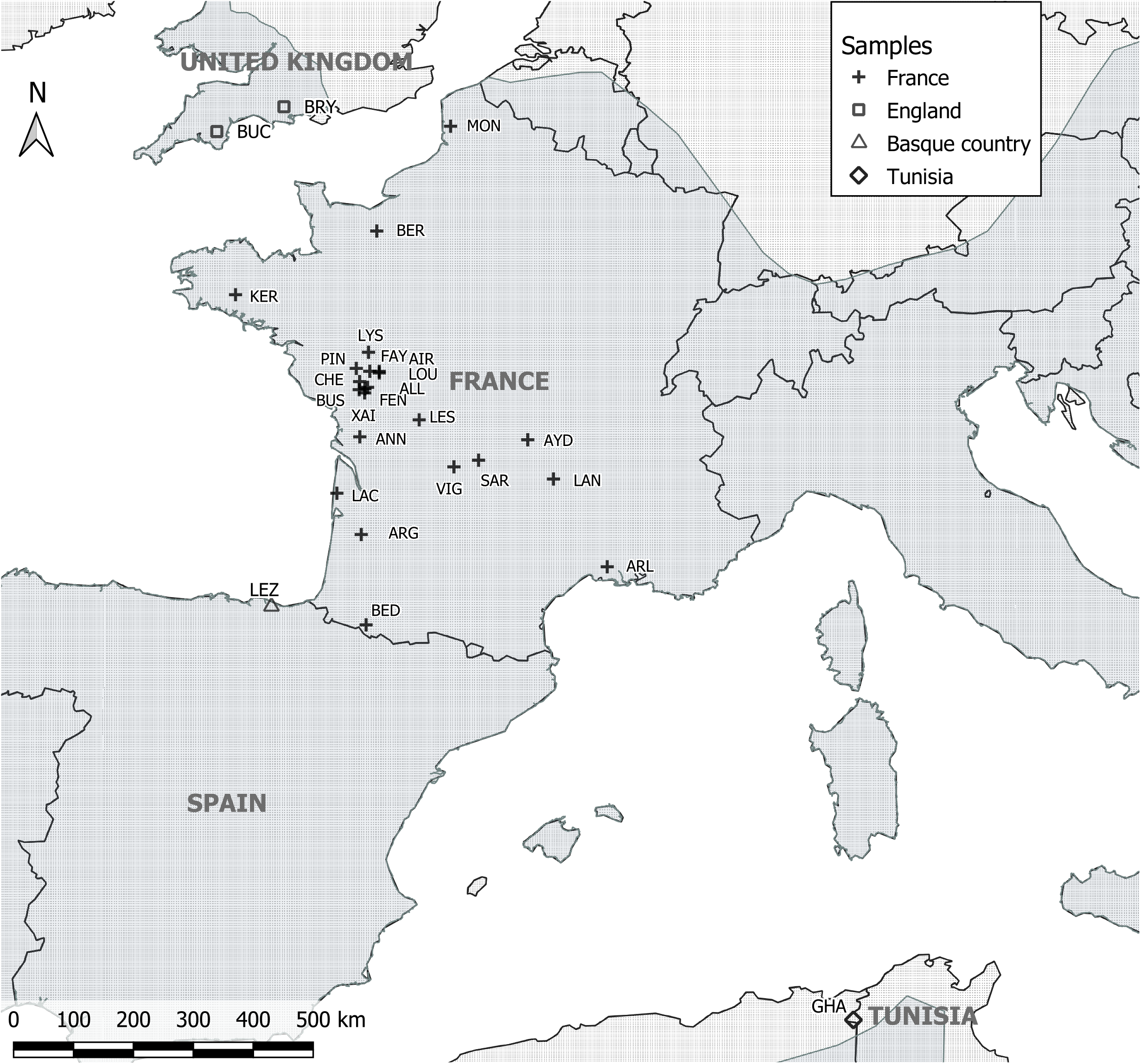
Map of the colonies included in this study. The geographic range of the species is colored in grey according to the IUCN data (2016). Bryanston (BRY), Buckfastleigh (BUC), Montreuil-sur-mer (MON), Kernascleden (KER), Lys-Haut-Layon (LYS), Allonne (ALL), Le Busseau (BUS), Le Pin (PIN), La Chapelle-Saint-Etienne (CHE), Xaintray (XAI), Airvault (AIR), Saint-Loup-sur-Thouet (LOU), Faye I’Abesse (FAY), Fenioux (FEN), Lessac (LES), Annepont (ANN), Sarran (SAR), Vignols (VIG), Aydat (AYD), Langeac (LAN), Lacanau (LAC), Argelouse (ARG), Arles (ARL), Bedous (BED), Lezate (LEZ), GHA.

**Table 1.**
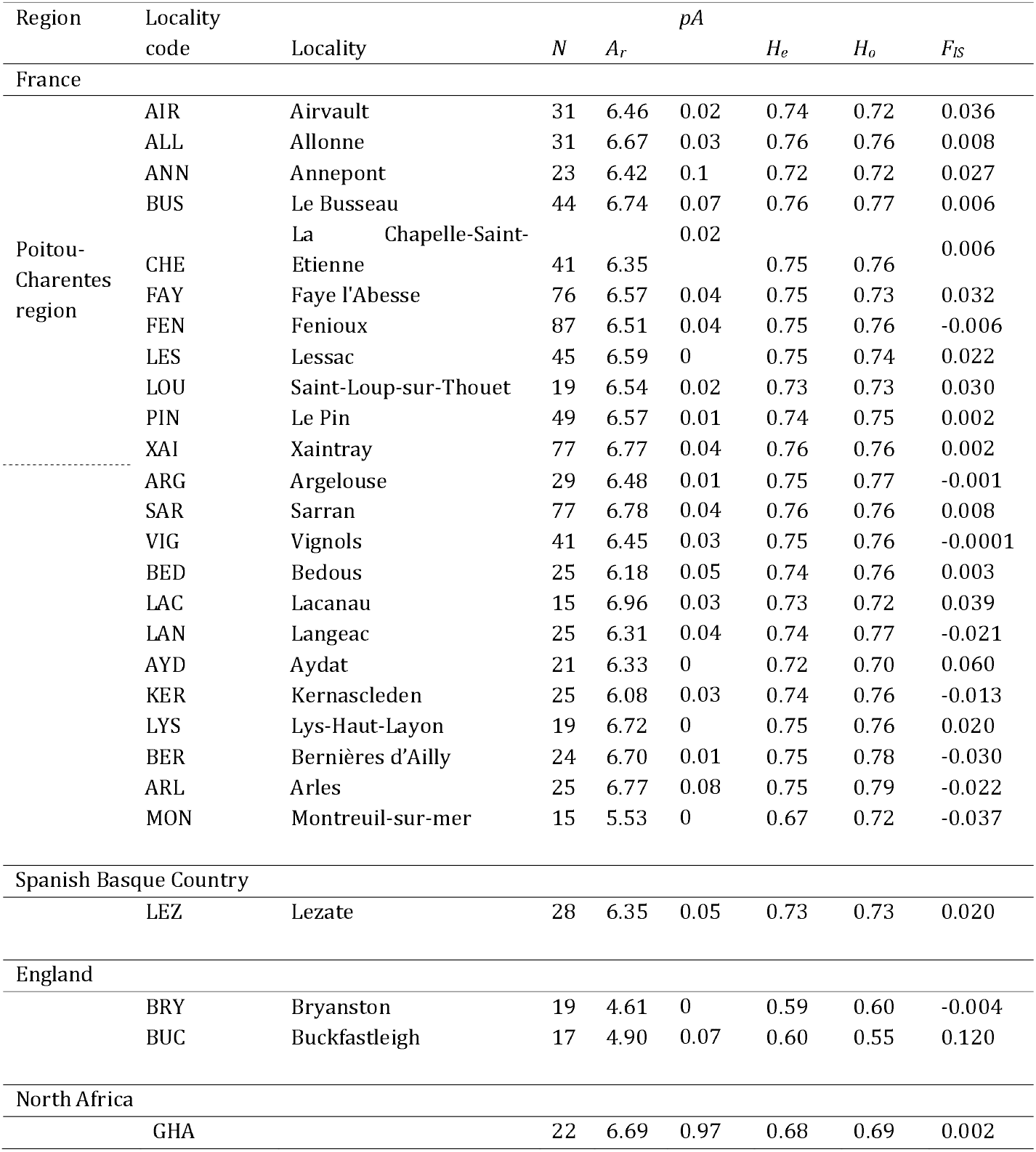
Genetic diversity parameters for each locality studied in France, Spanish Basque country, England and Tunisia. Sample size (N), corrected allelic richness (*A_r_*), corrected private allele richness (pA), expected (*H_e_*) and observed (*H*_0_) heterozygosity and inbreeding coefficient (*F_IS_*).

For each bat, a tissue sample was collected from the wing membrane (patagium) using a 3-mm diameter biopsy punch. Samples were preserved in 95° ethanol and quickly stored at 4°C until DNA extraction. Biopsy punches were cleaned with bleach, water then ethanol between each sample. Adults were distinguished from juveniles by trans-illumination of the cartilaginous epiphyseal plates in the phalanges (Anthony, 1988)

### DNA extraction and microsatellite genotyping

DNA was extracted from each wing sample using the EZ-10 Spin Column Genomic DNA Minipreps Kit for Animal (BioBasic) following the manufacturer’s instruction with a final elution of twice 50μL in elution buffer. We amplified 17 microsatellite loci using primers slightly modified from those previously designed for *R. ferrumequinum* (Dawson et al., 2004; Rossiter et al., 1999) and the lesser horseshoe bat (Puechmaille et al., 2005) (Table S1). Amplifications were organized into three multiplex PCRs that were conducted in 10μL reaction volumes containing 2μL of extracted DNA, 5μL of Multiplex Taq (Qiagen), 0.2μL of each primer (10μM), filled up with H2O to a volume of 3μL. We used the following PCR conditions: 95 °C for 15 min, 35 cycles of 94 °C for 30 s, 57 °C for 90 s, and 72 °C for 60 s, and a final elongation step at 60 °C for 30 min. 20μL of H20 were added to 10μL of each products in a new plate. 2μL of this final solution were mixed with 15μL of Formamide and 0.08μL of GeneScan 500 LIZ. The samples were genotyped using an ABI3130 automated sequencer and the GeneMapper v.5.0 software. Two independent readings were performed by different people to minimize genotyping errors.

Null alleles, large allele dropout, stuttering and scoring inconsistencies were tested for each colony using MICROCHECKER (Van Oosterhout, Hutchinson, Wills, & Shipley, 2004). Null allele frequencies were estimated for each locus and population using FreeNA, with 1,500 replicates for the computation of the bootstrap 95% confidence intervals (Chapuis & Estoup, 2007). We performed the exact tests implemented in GENEPOP v4.6 (Francois Rousset, 2008) to detect linkage disequilibrium for each pair of loci in each colony, and to investigate departure from Hardy–Weinberg equilibrium (HWE) for the whole dataset and for each colony and for each locus under the hypothesis of heterozygote deficit. We used the false discovery rate (FDR) to account for multiple testing (Benjamini & Hochberg, 1995).

### Genetic diversity and relatedness within colonies

We assessed genetic diversity within colony by estimating the allelic richness and private allele richness corrected for minimal sample size (*A_r_* and *pA*, N=13), the expected (*H_e_*) and observed (*H_o_*) heterozygosities, using respectively FSTAT v.2.9.3.2 (Goudet, 1995), HP-RARE (Kalinowski, 2005) and GENETIX v.4.05.2 (Belkhir, Borsa, Chikhi, Raufaste, & Bonhomme, 2004). Differences in *A_r_, H_e_* and *H_o_* between colonies were evaluated using Kruskall-Wallis tests in R 3.5.1 (R Core Team, 2016).

We estimated the fixation index *F_IS_* (Weir and Cockerham, 1984) using GENEPOP v4.6. We next estimated the maximum likelihood pairwise coefficient of relatedness *r* between pairs of individuals on an absolute scale (0, unrelated to 1, identical individuals), using the ML-RELATE software (Kalinowski, Wagner, & Taper, 2006). Indeed maximum likelihood estimate is less biased than commonly used estimators (Milligan, 2003). The coefficient *r* corresponds to the probability for each locus that individuals share zero, one or two alleles that are identical by descent. We used the genetic clusters identified from further clustering analyses (see below) as population references for estimating *r* within each colony.

### Population structure and genetic differentiation between colonies

The genetic differentiation between colonies was quantified using estimates of global and pairwise *F_ST_* (Weir and Cockerham, 1984). Significance was assessed using exact G-test of differentiation implemented in GENEPOP v4.6 (Rousset, 2008). Pairwise *F_ST_* and exact G-tests were computed for each pair of colonies and each pair of genetic clusters identified from further clustering analyses. We accounted for multiple testing using false discovery rate (FDR). In order to control for potential effects of null alleles on genetic differentiation, we also estimated pairwise *F_ST_* corrected for null-alleles using the ‘Excluding Null Alleles’ (ENA) correction implemented in FreeNA (Chapuis & Estoup, 2007).

Genetic structure was also investigated using the clustering approach implemented in the STRUCTURE program v2.3.4 (Pritchard, Stephens, & Donnelly, 2000). We determined the most likely number of genetic clusters using the log likelihood of *K* and *ΔK* statistic (Evanno, Regnaut, & Goudet, 2005) implemented in the website STRUCTURE HARVESTER (Earl & vonHoldt, 2012). We used the admixture model with uncorrelated frequencies and an alpha-value of 1 /*K*, as recommended by Wang (2017) in the case of unbalanced sampling. Same results were obtained with the per-default alpha-value and when applying or not the LOCPRIOR model to the population model (Hubisz et al. 2009). We performed 20 independent runs with a burn-in period of 1,000,000 iterations and 50,000 MCMC repetitions after burn-in, testing *K*= 1 to *K*=28 (N colonies + 1). We used the R package *pophelper* (Francis, 2017) to compute the plots. In addition we performed a Principal Component Analysis (PCA) implemented in the R packages *ade4* (Dray & Dufour, 2007) and *factoextra* (Kassambara & Mundt, 2017). Contrary to STRUCTURE algorithm, this approach does not rely on any specific population genetic assumption including Hardy-Weinberg equilibrium and linkage equilibrium (Pritchard et al., 2000).

We next used MAPI program (Mapping Averaged Pairwise Information, Piry et al., 2016), implemented in the R package *mapi*, to detect spatial genetic (dis-)continuity. This approach has low sensitivity to potential confounding effects resulting from isolation-by-distance (IBD) and does not require predefined population genetic model. It is based on a spatial network in which pairwise genetic distance between georeferenced samples are attributed to ellipses. A grid of hexagonal cells covers the study area and each cell receives the weighted arithmetic mean of the pairwise genetic distance associated to the ellipses intersecting the cell (Piry et al., 2016). We used the Rousse?s coefficient *â* (Rousset, 2000) computed with SPAGeDi 1.4 (Hardy & Vekemans, 2002) as an index of pairwise genetic differentiation between individuals and default parameter value for the eccentricity of the ellipses (0.975). We used the permutation procedure (1000 permutations) to identify areas exhibiting significantly higher or lower levels of genetic differentiation than expected by chance.

Lastly, isolation by distance (IBD) was analyzed with the regression of the genetic distances between colonies (*F_ST_* /1-*F_ST_*, Rousset, 1997) and the logarithm of the Euclidean geographic distances, implemented in GENEPOP v4.6. Confidence intervals and significance of regression slope and intercept were assessed by bootstrapping over loci. It was then tested using Mantel tests with 10,000 permutations. IBD analyses were performed on the complete dataset, within France and between pairs of colonies sampled in the different countries. The comparison of the results gathered from these IBD analyses should enable the identification of geographic barriers (Pyrenees, Channel and Mediterranean Sea) that might reduce *R. ferrumequinum* gene flow.

### Inference of demographic parameters

We first inferred the demographic history of *R. ferrumequinum* colonies. We used the software MIGRAINE v.0.5.1 (Leblois et al., 2014) and the model *OnePopVarSize.* This model assumes a unique variation in population size of an isolated panmictic population starting *T* generations ago, that was followed by continuous exponential change in population size until the moment of sampling. This model estimates three scaled parameters: the current scaled population size (*ϑ* = 4*N_e_μ*) and the ancestral scaled population size (*ϑ_anc_* = 4*N_anc_μ*), where *N_e_* and *N_anc_* are the current and ancestral diploid population sizes and *μ* is the mutation rate per generation of the loci, and *D*, which is the time when the demographic change started scaled by current population size (*D*=*T*/4*N*). The mutation model used was a generalized stepwise mutation model (GSM) which is characterized by a geometric distribution of mutation steps with the parameter pGSM. We used the ratio *Nratio* (*N/N_anc_*) to characterize the strength of demographic events. In case of a contraction, the *Nratio* is < 1 and in a case of an expansion it is > 1. The change in population size was considered significant when the 95% confidence intervals (95% CIs) of the *Nratio* did not include the value “1”. Runs were performed independently for each colony and for each genetic cluster that was previously identified. We used 1,000 to 20,000 trees per points, 600 to 800 points and eight iterations by run. When no signal of demographic change was found, we used the *OnePop* model implemented in MIGRAINE to estimate the scaled effective population size *N_e_* (*ϑ*= 4*N_e_μ*) of stable and panmictic population.

We then inferred contemporary levels and directions of migration between the main genetic clusters using the program BAYESASS v3.0.4 (Wilson & Rannala, 2003). We performed 5 independent runs of 10,000,000 iterations sampled every 2000 iterations, with a burn-in of 1,000,000. For each run we calculated the Bayesian deviance using the R script provided by Meirmans (2014). We used this deviance as a criterion to find the run that provided the best fit and to identify runs with convergence problems (Faubet, Waples, & Gaggiotti, 2007; Meirmans, 2014). We ran preliminary runs to adjust the maximum parameter change per iteration (Delta values). It is important to optimize the acceptance rates for proposed changes to parameters (20% to 60% is ideal; Wilson and Rannala, 2003). The adjustments are important because if the acceptance rates are too low or too high the chain does not mix well and it fails to adequately explore the state space. In the final run, we used delta values of respectively 0.45, 0.40, and 0.51 for allele frequency, migration and inbreeding.

## Results

Three of the 17 microsatellites genotyped were excluded from further genetic analyses: one of them was monomorphic and the two others had poor quality profiles. Results gathered using MICROCHECKER showed no large allele dropout, neither scoring inconsistencies due to stuttering. Null alleles were suspected at loci Rferr06 in three colonies from France (‘AIR’, freq = 0.074; ‘FEN’, freq= 0.046; ‘XA’, freq = 0.041), at loci RHD103 (France, ‘ARL’, freq = 0.078), Rferr27 (France, ‘BED’, freq = 0.152) and Rferr01 (England, ‘BUC’, freq = 0.149). Nevertheless we did not detect any deviation from HWE in any colony. Fifteen out of the 2418 pairs of loci (0.62%) exhibited significant linkage disequilibrium but the loci involved were not consistent among colonies. Therefore, we did not exclude any other loci.

### Genetic diversity and relatedness

A summary of the genetic diversity indices (*F_IS_, A_r_, H_e_, H_o_*) based on the 14 validated microsatellites is presented in Table 1. Global Hardy-Weinberg test (H1 = heterozygote deficit) was significant (*p* = 0.017, *F_IS_* = 0.009). Local *F_IS_* estimates (*i.e.* for each colony) ranged from −0.037 (‘MON’, Northern France) to 0.120 (‘BUC’, England). All tests of departure from local HWE were non-significant after FDR correction (*p* > 0.05). We did not find significant differences of genetic diversity between colonies (Kruskal-Wallis tests, X^2^_*Ar*_ = 18.638, *df_Ar_* = 26, *p_Ar_* = 0.851 and *X*^2^_*He*_ = 26, *df_He_* = 26, *p_He_* = 0.463), although the English colonies, the Northern French colony ‘MON’ and the Tunisian colony ‘GHA’ showed lower value of allelic richness and expected heterozygosity than the others (Table 1, Figure 2). Private allele richness was low in all colonies except the Tunisian one (‘GHA’; Table 1).

**Figure 2.**
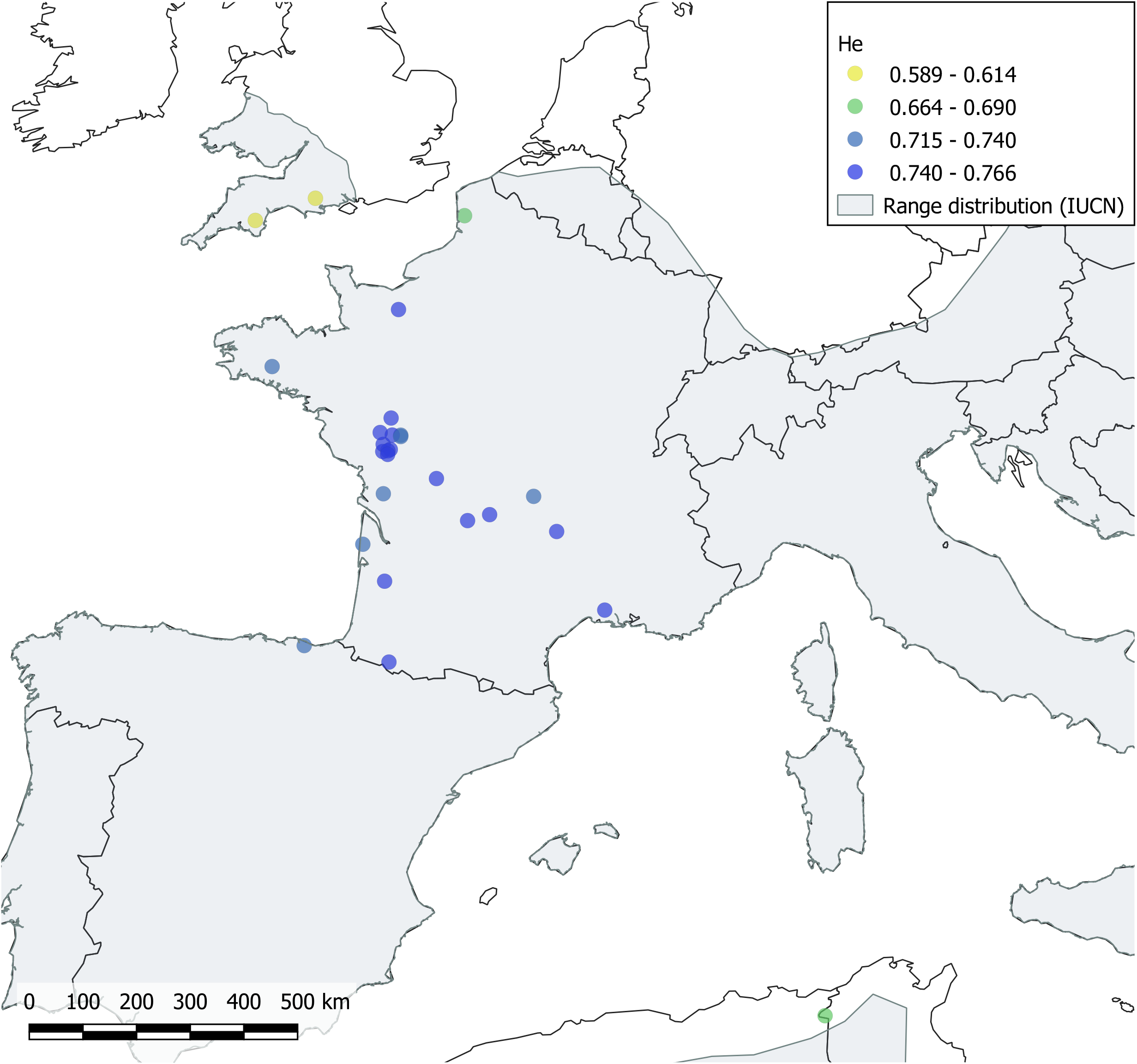
Expected heterozygosity (*H_e_*) estimated for each colony. Yellow to blue colors indicate low to high levels of *H_e_*. The geographic range of *R. ferrumequinum* is shaded in grey.

The distributions of the pairwise coefficient of relatedness *r* within each colony showed a common pattern in all colonies except in the Northern French colony ‘MON’ (Figure 3). For all colonies, the distribution of the *r* coefficient was L-shaped with a peak of unrelated individuals (*r* = 0) and a decreasing proportion of related individuals. In the Northern French colony ‘MON’, we observed a more uniform proportion of unrelated and relatively closely related individuals (*r* ranging between 0.0 and 0.3). When considering pairwise relatedness between individuals from different colonies, we still observed high levels of relatedness (*r* > 0.5) (Figure 4). The high levels of relatedness involved female-female pairs and female-male pairs but never male-male pairs. It concerned 537 females, 33 males and one undetermined individual. The majority of the females were adults (474 adults, 59 juveniles and four undetermined), what was not the case when considering males (12 adults, 19 juveniles and two undetermined).

**Figure 3.**
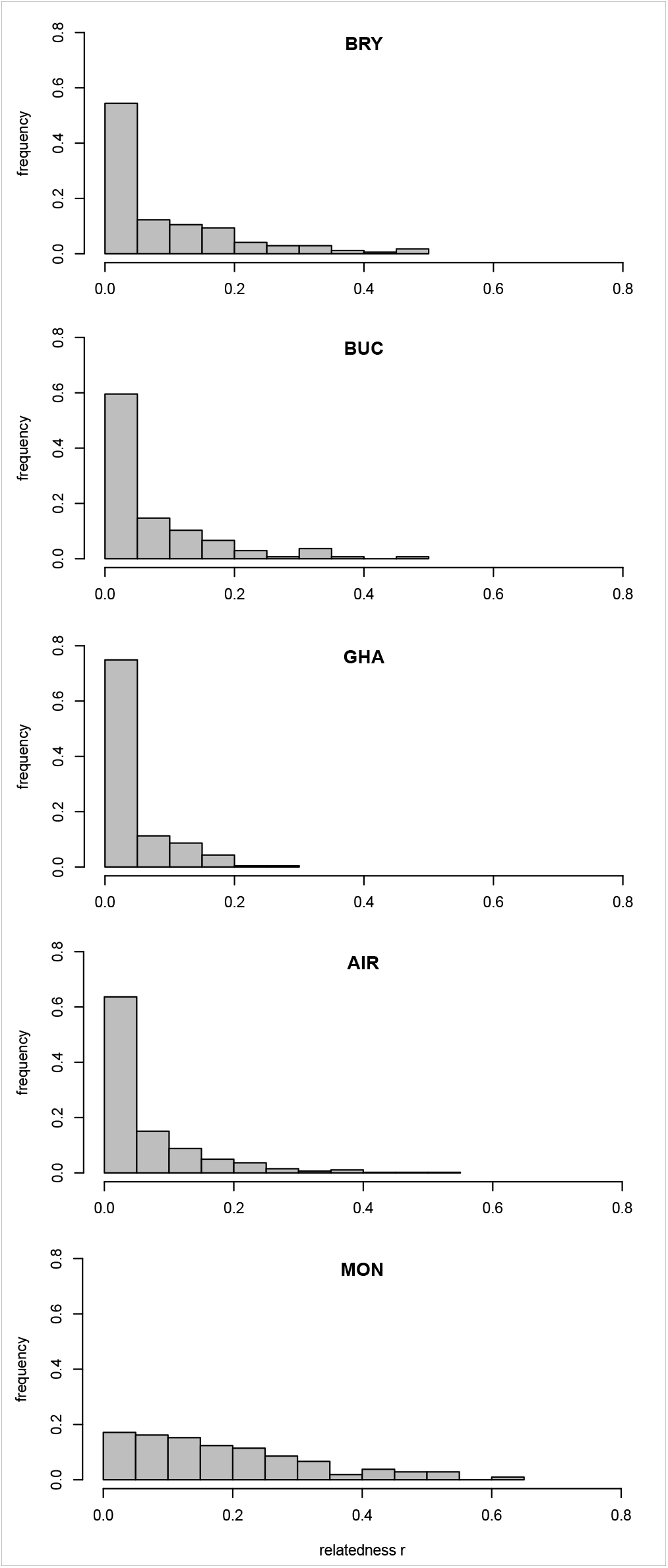
Distribution of the pairwise relatedness coefficient r estimated with the ML-relate software (Kalinowski et al., 2006) within colonies from France (AIR, MON), England (BRY, BUC) and Tunisia (GHA).

**Figure 4.**
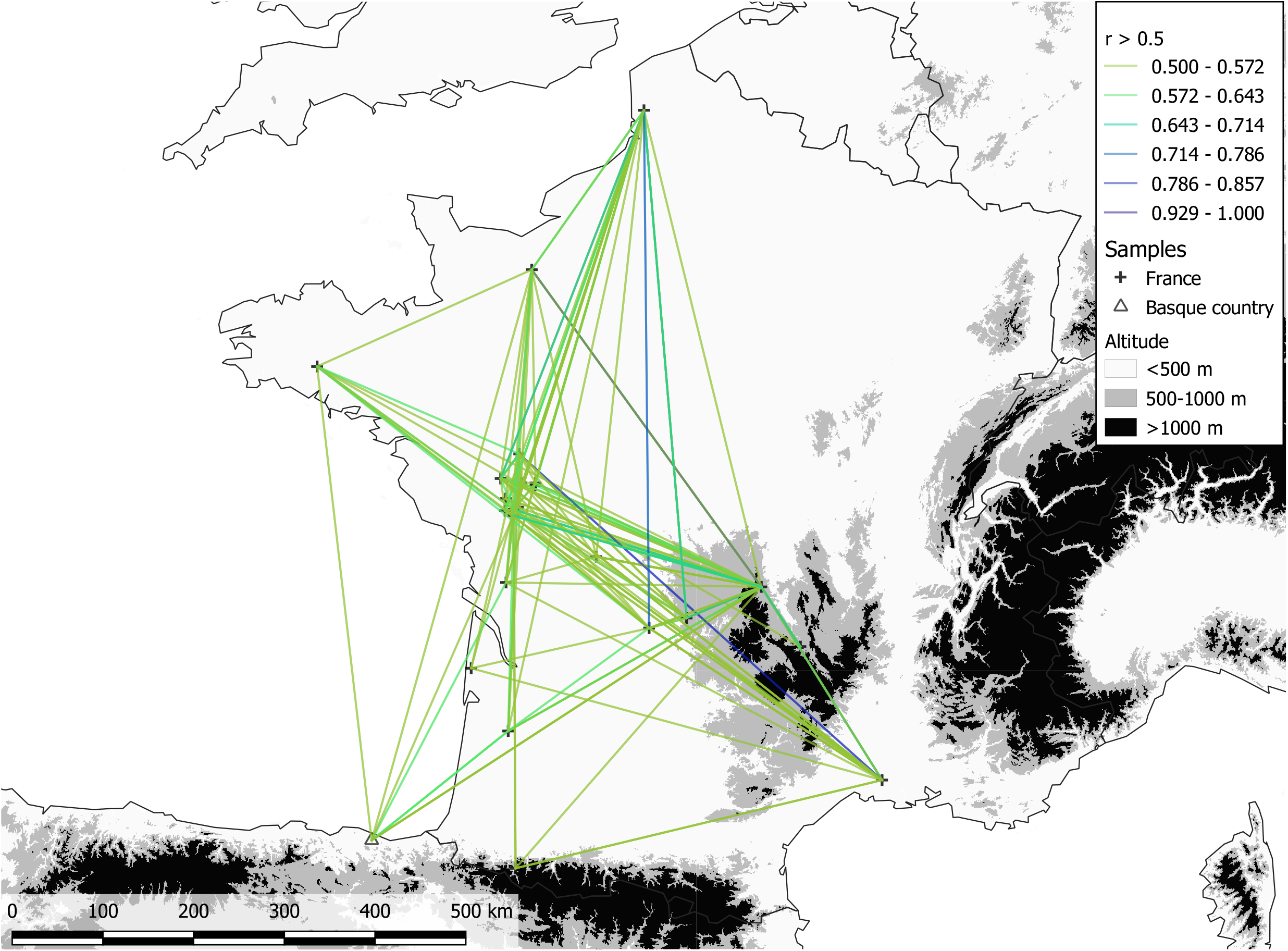
Geographic distribution of the higher values of pairwise relatedness coefficient (*r* > 0.5) between *R. ferrumequinum* colonies from the continental genetic cluster (France and the Spanish Basque Country). Yellow to blue colors indicate medium to high levels of *r*.

### Population structure and genetic differentiation between colonies

The pairwise *F_ST_* estimates between colonies and associated G-tests are presented in Supplementary Table 2. *F_ST_* estimates calculated with and without excluding null alleles (ENA) were very similar; therefore we only reported the uncorrected estimates. Low but significant genetic differentiation was observed between colonies within France (*F_ST_* < 3%, 65.24% of the G-tests with *p* < 0.05). The Northern French colony ‘MON’ exhibited higher estimates of pairwise *F_ST_* than the other colonies (3.49% < *F_ST_* < 6.46%, G-tests *p* < 10^-3^). Among the French colonies, ‘MON’ was also the colony that exhibited the highest levels of genetic differentiation with the English (*F_ST_* > 16%, G-tests *p* < 0.001), Spanish Basque (*F_ST_* = 7.17%, G-tests *p* < 0.001) and Tunisian colonies (*F_ST_* = 17.05%, G-tests *p* < 0.001). The Spanish Basque colony ‘LEZ’ was more genetically differentiated from colonies in England and Tunisia (England: 9.88% < *F_ST_* < 11.9%, G-tests *p* < 0.001 and Tunisia: *F_ST_* = 14.87 %, G-tests *p* < 0.001) than from colonies in France (*F_ST_* < 3%).

Using STRUCTURE, we found that the *ΔK* statistics (Evanno et al., 2005) were highest for *K*=2 but the likelihood of the number of genetic clusters Ln (L(*K*)) showed similar values for *K*=2 and *K*=3 (Figure S1). For *K*=2, the first genetic cluster included all colonies from England, France and Spanish Basque Country and the second genetic cluster included the Tunisian colony. For *K=*3, the first main genetic cluster included all colonies from the European mainland. The second main genetic cluster included the two colonies from England and the third genetic cluster included the Tunisian colony (Figure 5). Increasing *K* did not change this clustering pattern. The PCA showed similar results, with three main genetic clusters (England, European mainland and Tunisia, Figure S2).

**Figure 5.**
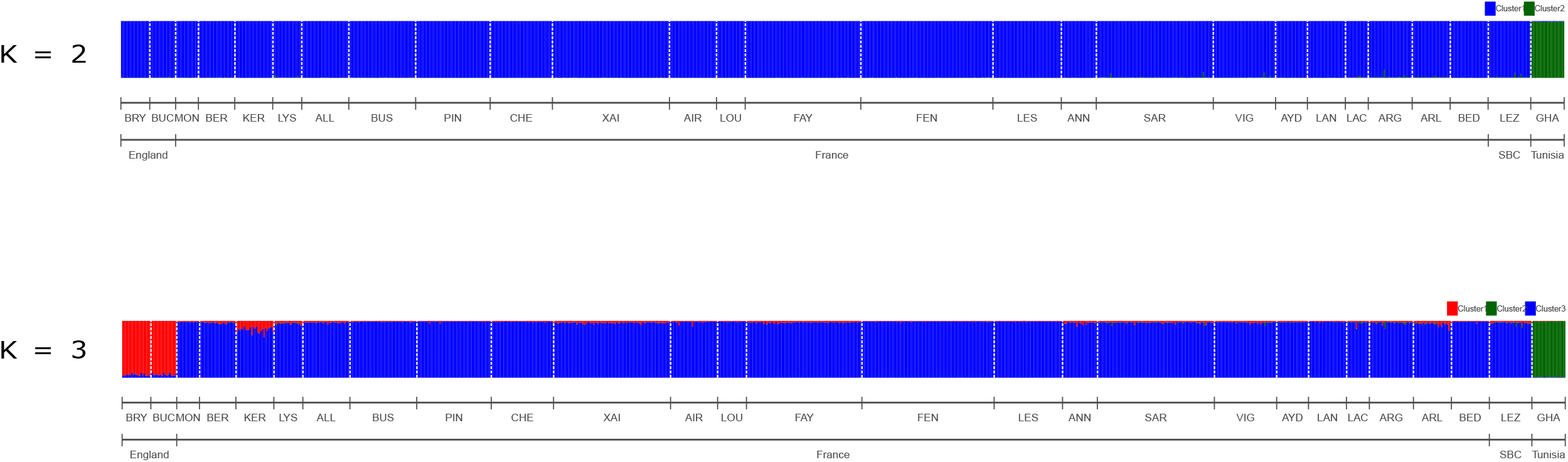
STRUCTURE plots for K=2 and K=3 clusters. Each bar represents an individual colored according to its membership probability to a given cluster. Individuals are sorted by localities. The white dashed lines separate the 27 colonies where the individuals were sampled: Bryanston (BRY), Buckfastleigh (BUC), Montreuil-sur-mer (MON), Kernascleden (KER), Lys-Haut-Layon (LYS), Allonne (ALL), Le Busseau (BUS), Le Pin (PIN), La Chapelle-Saint-Etienne (CHE), Xaintray (XAI), Airvault (AIR), Saint-Loup-sur-Thouet (LOU), Faye I’Abesse (FAY), Fenioux (FEN), Lessac (LES), Annepont (ANN), Sarran (SAR), Vignols (VIG), Aydat (AYD), Langeac (LAN), Lacanau (LAC), Argelouse (ARG), Arles (ARL), Bedous (BED), Lezate (LEZ) and GHA. SBC = Spanish Basque Country.

While including all colonies, the program MAPI revealed that the Mediterranean Sea corresponded to an area of significantly higher genetic dissimilarity (Figure 6A). A second analysis excluding the Tunisian colony highlighted a significantly higher genetic dissimilarity between the English and the European mainland colonies, and a genetic homogeneity on the continent (Figure 6B).

**Figure 6.**
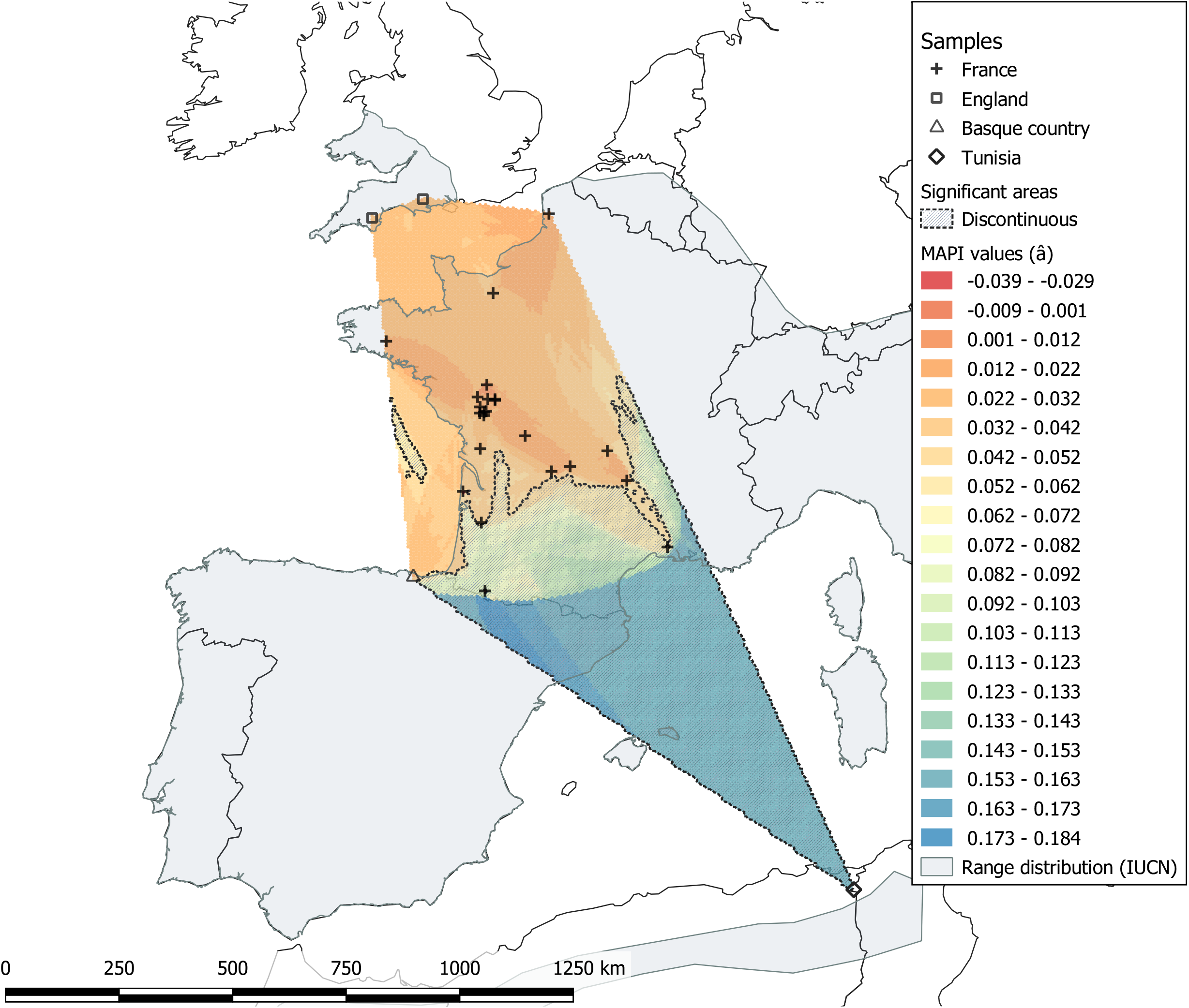
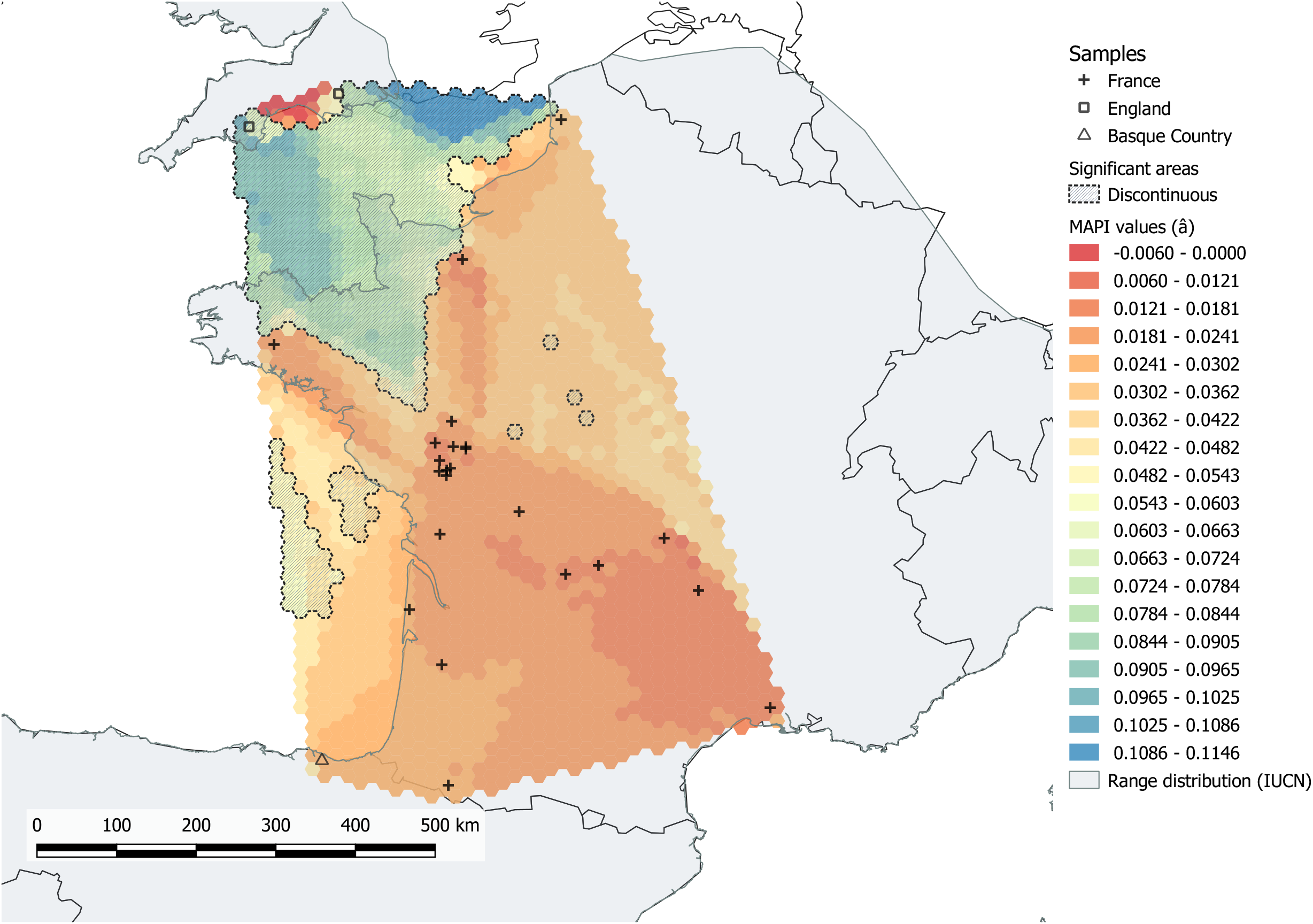
Geographic distribution of the pairwise Rousse’s genetic distance â (Rousset, 2000) resulting from MAPI (Piry et al., 2016). A) All colonies are included. B) The Tunisian colony is not included in the analysis. Dots correspond to the colonies sampled in this study. Levels of genetic dissimilarity are indicated using a color scale ranging from red (lower genetic dissimilarity) to blue (higher genetic dissimilarity). Significant areas are represented by dashed lines.

We found a significant positive relationship between genetic differentiation and geographic distance within France (slope of the regression = 0.0063 [0.0042 – 0.0090]; *p_Mantel_* = 0), between France and Spanish Basque Country (slope of the regression = 0.0212 [0.0120, 0.0412]; *p_Mantel_* = 10’^4^) and between France and Tunisia (slope of the regression = 0.0457 [0.0009, 0.1390]; *p_Mantel_* = 10”^3^). The Northern French colony ‘MON’ was more genetically differentiated from the other French colonies (Figure 7). When excluding this particular colony ‘MON’ from the regression analyses, we still found a positive relationship between genetic differentiation and geographic distance among French colonies but the slope was weaker (slope of the regression = 0.0025 [0.0015 – 0.0036]; *p_Mantel_* = 0). Moreover, considering only differentiation between pairs for French-English, French-Basque and French-Tunisian colonies without including the Northern French colony ‘MON’, the isolation by distance relationship was not significant (Table 3). All results are presented in Table 3 and Figure 7.

**Figure 7.**
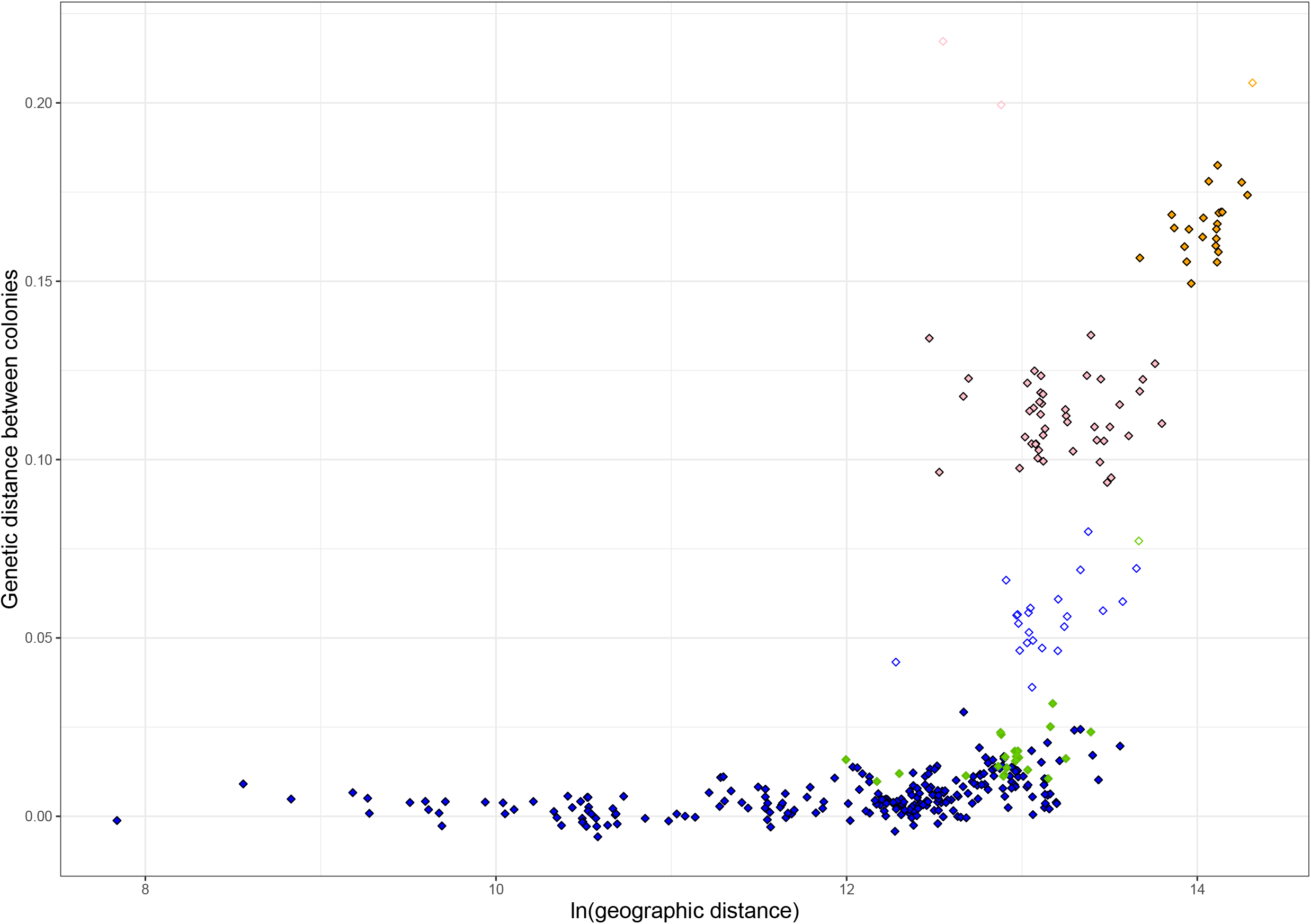
Plots of the within/between isolation by distance analyses. The genetic distance between colonies was estimated as (*F_ST_* /1-*F_ST_*)· The color of the dots indicate the geographic origin of the pairs of colonies: within France (blue), between France and Spanish Basque Country (green), between France and Tunisia (orange) and between France and England (pink). The pairs of colonies that include the French colony ‘MON’ are not color filled.

**Table 2.**
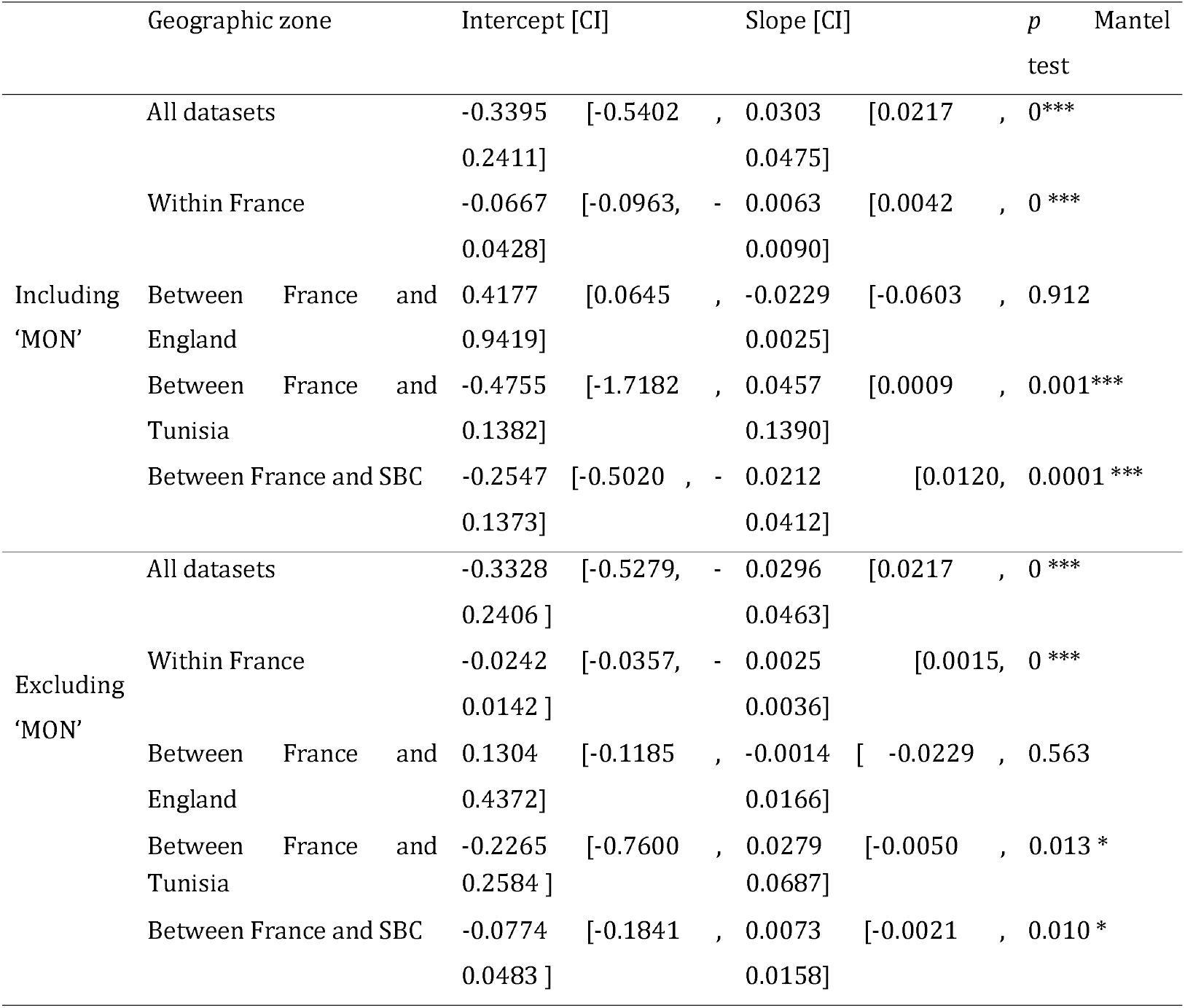
Isolation by distance characteristics using the genetic differentiation parameter *(F_ST_* /1-*F_ST_*) between colonies against the logarithm of the Euclidian geographical distance. 95% confidence intervals (Cl) for the slope and the intercept of the IBD were obtained by ABC bootstrapping. Significant p-values of Mantel tests are represented in bold with *** for *p* < 0.001, ** for *p* < 0.01 and * for *p* < 0.05. ‘MON’ is the Northern French colony of the study area, SBC = Spanish Basque Country.

**Table 3.** Means of the posterior distributions of contemporary migration rate *(m)* with standard deviation in parentheses. Migration and non-migration rates significantly greater than 0 are represented in bold.

| Migration into | Migration from |  |  |  |  |
| --- | --- | --- | --- | --- | --- |
|  | Genetic cluster 1 |  |  | Genetic cluster 2 | Genetic cluster 3 |
|  | MON | France | Basque | England | Tunisia |
| MON | **0.6825**<br>**(0.0152)** | **0.2700**<br>**(0.0276)** | 0.0161<br>(0.0153) | 0.0158<br>(0.0148) | 0.0156<br>(0.0153) |
| France | 0.0004<br>(0.0004) | **0.9975**<br>**(0.0013)** | 0.0004<br>(0.0004) | 0.0012<br>(0.0011) | 0.0004<br>(0.0004) |
| Basque | 0.0101<br>(0.0095) | **0.2921**<br>**(0.0192)** | **0.6768**<br>**(0.0101)** | 0.0109<br>(0.0108) | 0.0102<br>(0.0100) |
| England | 0.0082<br>(0.0081) | 0.0128<br>(0.0116) | 0.0083<br>(0.0082) | **0.9626**<br>**(0.0169)** | 0.0080<br>(0.0078) |
| Tunisia | 0.0129<br>(0.0125) | 0.0126<br>(0.0121) | 0.0129<br>(0.0122) | 0.0124<br>(0.0119) | **0.9493**<br>**(0.0229)** |

### Inference of demographic parameters

Using MIGRAINE, we detected a significant signature of expansion for the Tunisian colony (‘GHA’; *N_ratio_* = 2989 [1.63 – 8063]), with an estimated origin between one and 11,000 generations (D = 0.811 [2.17 e-06 – 2.201]). None of the other colony or genetic cluster showed significant signature of demographic change (Table S3). The marginally significant signature of contraction observed for the Western French colony ‘CHE’ in the Poitou-Charentes region was also considered to be nonsignificant because the higher value of the confidence interval was very close to 1 (0.99) and because we performed a high number of tests.

We estimated the scaled current population size *ϑ* (4*N_e_μ*) for all stable colonies, *ϑ* estimates ranged between 1.854 and 6.465 (Table S4, Figure 8). The lowest estimates were found for the English colonies (‘BRY’ *ϑ* = 1.854; ‘BUC’ *ϑ* =2.107) and for the Northern French colony ‘MON’ (*ϑ* = 3.030). Other colonies and pool of colonies exhibited similar levels of *ϑ* estimates (from 4.086 to 6.465). Kruskal-Wallis tests and post-hoc pairwise comparisons of *ϑ* estimates using Wilcoxon test (with FDR correction for multiple testing) revealed a significant difference of *ϑ* estimates between the European mainland and the English genetic clusters (*p* = 0.002) and non-significant differences between the other clusters (*p* > 0.115). We observed a significant negative relationship between the estimated *ϑ* of each colony and the distance of the colony to the centroid of our sampling (*p* < 0.05; Figure S3).

**Figure 8.**
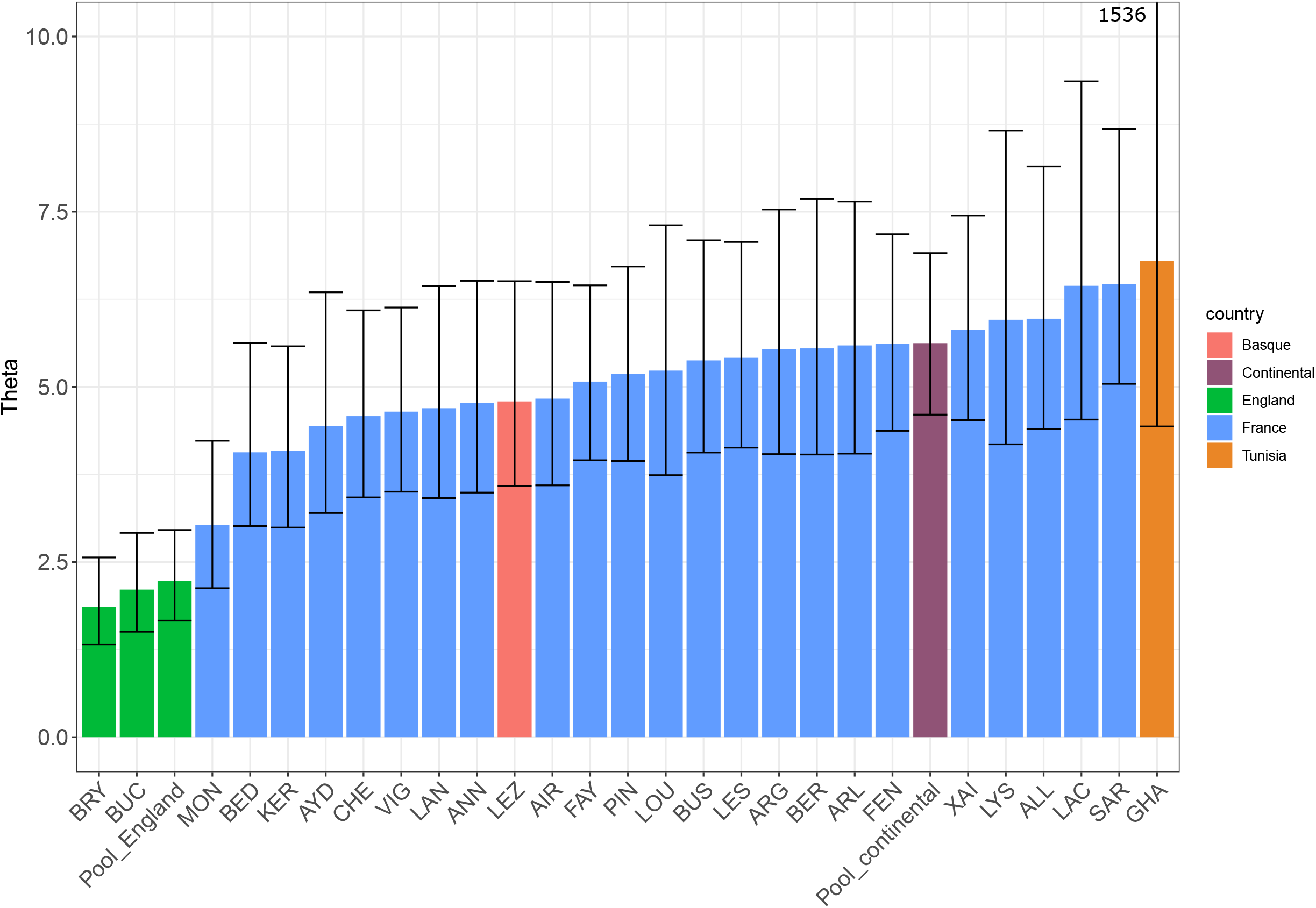
Estimation of *ϑ* (4*N_e_μ*) for each colony and genetic cluster based on the OnePop model for stable colonies and OnePopVarSize for the Tunisian colony ‘GHA’ which experienced expansion. As the confidence interval of Tunisian *ϑ* is very large (4.435 – 1536), the superior born was not represented to avoid the flattening of the graph. Pool_England and Pool_continental respectively correspond to the English and French-Spanish Basque colonies. Colonies are classified by *õ* estimates.

The five repeats runs implemented in BAYESASS provided similar results and converged well, with values ranging from *D_run2_* = 67128.6 to *D_run4_* = 67134.7. None of the migration rates estimated between the three main genetic clusters (England, European mainland, Tunisia) were significant (Table 4). Within the European mainland cluster we found significant unidirectional migration rates from the French to Spanish Basque colonies (*m* = 0.2921 ± 0.0192). As our results emphasised particular signatures for the Northern French colony ‘MON’ all along the study, we also estimated migration rates considering ‘MON’ apart from the other French colonies. Our results revealed significant unidirectional migration rates from the French colonies to the Northern French colony ‘MON’ (*m* = 0.2700 ± 0.0276).

## Discussion

### A large and stable population of *R. ferrumequinum* in Western European mainland

Our study revealed high and homogeneous levels of genetic diversity within the European mainland colonies examined. These levels were similar to those previously detected in colonies from the eastern parts of *R. ferrumequinum* distribution (Rossiter et al., 2007; Rossiter et al., 2000), or in other bat species such as *Rhinolophus euryale* and *Myotis myotis* (Budinski et al., 2019; Castella, Ruedi, & Excoffier, 2001). We did not find evidence of departure from Hardy-Weinberg equilibrium within colonies or when considering the European mainland colonies altogether. We also found very low levels of genetic differentiation between these European mainland colonies of *R. ferrumequinum*, as indicated by *F_ST_* estimates. These colonies were identified as a unique genetic cluster according to the results provided by STRUCTURE and MAPI analyses and the slight isolation by distance pattern detected. Altogether, these results indicated high gene flow between colonies at this scale and suggested that these European mainland colonies form a single, large and panmictic population. Therefore, our results supported the previous studies of *R. ferrumequinum* phylogeography which had revealed a unique genetic cluster in Western Europe (Flanders et al. 2009; Rossiter et al., 2007). In complement of these findings, we could infer the effective size *ϑ* (4*N_e_μ*) of this large population. We found homogeneous *ϑ* among the European mainland colonies and for the pool of these colonies. This result was congruent with our finding of one large population with each colony representing a replicate of the whole population. It is not trivial to transform this estimate *ϑ* into an effective number of individuals (*N_e_*) as it requires fixing the mutation rate of microsatellite markers in this species. This is a crucial issue because estimates of mutation rates can greatly vary between and within species, and any inaccuracy in the mutation rate estimate is propagated in the estimation of *N_e_* (Waples, 2010). As the mutation rate of *R. ferrumequinum* is not known, we were not able to precisely estimate its effective population size. However, based on the mutation rate range commonly used for microsatellites (10^-4^ to 10^-3^), our analysis supported a minimum effective population size of approximatively 1316 [1150 – 1728] individuals and a maximum effective population size of approximatively 13,160 [11,508 – 17,273] individuals. Converting this effective population size *(N_e_)* in a number of adult individuals in the population *(N_c_:* Luikart et al., 2010) would be even more relevant for *R. ferrumequinum* management. The average ratio *N_e_/N_c_* = 0.34 is commonly used (Frankham, 1995), what would indicate that the number of adult individuals in the Western European mainland population could range between 3870 and 38,705. These estimates are lower than the 47,651 individuals estimated from previous ecological winter surveys conducted in France from 2001 to 2012 (Vincent and Bat Group SFEPM, 2014). But these values have to be taken very cautiously. On the one hand, the individual count included both adults and yearly juveniles. On the other hand it is known that providing estimates of *N_e_/N_c_* ratio also suffers from several important methodological issues (see Luikart et al., 2010 and Vonhof and Russell, 2015). For example, this average estimate of *N_e_/N_c_* is derived from species that may potentially have very different life histories compared to bats; this ratio may fluctuate and reductions in *N_e_* can occur without any impact on *N_c_*.

We did not reveal any signature of demographic decline, neither for the colonies of the French Poitou-Charentes region, nor for the delineated European mainland population of *R. ferrumequinum.* A stable population in the Poitou-Charente region was not expected as previous surveys of hibernation roost counts indicated a 30% population decline 15 years ago. Several explanations may underlie these discrepancies. First, Leblois et al. (2014) showed that the capacity to detect demographic events from genetic data depends on the number of genetic markers used, the strength of the event and the time when it happened. Therefore, because the decline was recent and we used 14 microsatellites, our study may have suffered from a lack of power that could explain the absence of signature of demographic decline in our data. Second, it is also likely that count-based analyses might have led to wrong estimates of the bat population demography, either because of a lack of standardized procedures through time or because knowledge of bat roosts was not exhaustive (Kunz et al., 2009; O’Shea et al., 2003). As such, previous studies based on ringing data have shown that the decrease in abundance observed at a given time may rather reflect punctual movements between hibernation roosts than local declines (Brosset & Poillet, 1985). Therefore, we emphasize the urgent need to more deeply investigate the demographic trends of *R. ferrumequinum* colonies within the Western European part of its range, using a combination of standardized joint genetic and ecological surveys.

### Dispersal and reproduction of *R. ferrumequinum*

Our results revealed that the Mediterranean Sea and the English Channel may act as barriers to *R. ferrumequinum* gene flow. Seas have often been shown to limit bat gene flow, irrespectively of bat flight capacity (García-Mudarra et al., 2009). The high genetic differentiation of the English colonies observed in this study was expected because of the potential founder effects associated with *R. ferrumequinum* colonization since the Late Glacial Maximum (LGM) (Flanders et al., 2009; Rossiter et al., 2007). Our demographic inferences also revealed the absence of current gene flow between the English and European mainland colonies. The high genetic differentiation observed between Tunisian and European mainland colonies may also rely on historical colonization history. *R. ferrumequinum* seems to have been present in North Africa before the Late Glacial Maximum (Flanders et al., 2009 ? Rossiter et al., 2007), and there is no evidence that North Africa was recolonized from Europe post-LGM. These historical patterns of genetic differentiation might have been reinforced by the current absence of gene flow between Europe and North Africa. However, we cannot exclude the possibility that the Strait of Gibraltar, which leaves a gap of just 14 km, connects populations of *R. ferrumequinum* across the Mediterranean Sea. The degree of permeability of the Mediterranean Sea for this bat species therefore deserves a dedicated study, aiming at assessing the potential pathways of *R. ferrumequinum* movements and gene flow between Europe and North Africa.

Similarly, our results regarding the Pyrenees end up with several alternatives. We did not find any evidence of gene flow disruption between both sides of the Western Pyrenees, suggesting that this mountain does not limit dispersal. On the one hand, this possibility may sound surprising as mountains have previously been shown to act as barriers to gene flow for several bat species (Dool et al., 2013; Razgour et al., 2013; Wright et al., 2018). Moreover, *R. ferrumequinum* is considered to be confined to lowlands (<1000m elevation; Le Roux et al., 2017) and previous genetic studies suggested that mountains such as the Alps may be a geographical barrier to this bat species (Flanders et al., 2009; Rossiter et al., 2007). On the other hand, the possibility that the Pyrenees do not prevent dispersal from both sides of this mountain may find persuasive explanations. Flanders et al. (2009) did not find evidence of gene flow disruption between the Pyrenees in a previous phylogeographic study. An explanation would be that *R. ferrumequinum* uses the shoreline as corridor along both the Atlantic and Mediterranean edges of the Pyrenees, where altitude is lower than 500m, as has been seen for migratory birds (Galarza & Tellería, 2003). Future genetic and ecological studies including dense sampling from both sides and all along the Pyrenees are now required to assess whether *R. ferrumequinum* movement and genetic mixing is restricted by mountains.

In the absence of important landscape barriers such as seas or, potentially, mountains, we showed that *R. ferrumequinum* is able to move over hundreds of kilometers, as exemplified by the low levels of genetic differentiation observed at large geographical scales, the inference of significant migration rates between European mainland maternity colonies and the high levels of relatedness observed between individuals sampled in distant colonies.

These results also revealed the high genetic mixing that occurs at large scale between *R. ferrumequinum* European mainland colonies. Several demographic processes may underlie this genetic mixing. First, mating dispersal at large distance would lead to extra-colony copulations and to the relaxation of colonies’ genetic borders (Veith, Beer, Kiefer, Johannesen, & Seitz, 2004). Second, because we also found high levels of relatedness between juveniles (under two-years old) sampled at considerable distances (up to 861km), natal dispersal (one-way movement of juveniles during their first year, from their colony of birth to another) and/or movements of adults from one maternity colony to another between two consecutive years reproduction events are also potential mechanisms shaping genetic mixing. These alternatives are still difficult to evaluate due to a lack of knowledge with regard to *R. ferrumequinum* mating behaviour (where and when) and dispersal in particular the one of males. In the future, long-term capture-mark-recapture surveys of adults and juveniles could provide invaluable information to assess the relative importance of mating, natal and breeding dispersal in the genetic mixing of colonies within management units.

### Differences in the functioning of central – peripheral and island – continental colonies

Our results revealed contrasting patterns of within and between population genetic structure when comparing *R. ferrumequinum* European mainland population with colonies from Tunisia, England and Northern France (‘MON’). In these latter colonies, we detected lower levels of genetic diversity (*H_e_* up to 20% lower) and smaller estimates of effective population size (up to half the size) than in the European mainland population. They were also more genetically differentiated than the European mainland ones, as revealed by *F_ST_* estimates, isolation by distance and clustering analyses. Some of these particular colonies might be insular (England), but common to all of them is to be located near the edge of *R. ferrumequinum* distribution range. Contrasting levels of genetic diversity between insular and continental populations are common in animals (Frankham, 1996; Frankham, 1995) and have already been observed between England and the continent in several bat species such as *R. ferrumequinum* (Rossiter et al., 2000), *Myotis bechsteinii* (Wright et al., 2018), *R. hipposideros* (Dool et al., 2013) and *Eptesicus serotinus* (Moussy et al., 2015). The contemporary isolation of colonies from UK with the continent might have maintained higher levels of genetic drift, as shown by the low *N_e_* estimates, which reinforces their vulnerability (Newman & Pilson, 1997). These colonies may face stochastic reduction of genetic diversity that could limit their evolutionary potentials, in particular in the face of environmental changes.

More surprisingly, the Northern French colony at Montreuil-sur-Mer (‘MON’) exhibited genetic patterns that were similar to those observed in the English colonies (lower level of genetic diversity, smaller effective population size and higher levels of genetic differentiation compared to the other European mainland colonies). This colony was also the only one that exhibited a high proportion of strongly related individuals. Overall, these results suggested that this colony experienced strong genetic drift, due to a small effective population size and a limited gene flow with other European mainland colonies. This situation could be explained by a lack of favourable habitats surrounding the colony, leading to limited dispersal and reduced extra-colony mating. It could also be explained by the location of the colony, at the northern limit of *R. ferrumequinum* distribution range (central-margin hypothesis, Eckert et al., 2008). Indeed, we observed asymmetric gene flow from the core to the edge of the European mainland population. We also detected a decrease of effective population size and genetic diversity and an increase of genetic differentiation while moving further away from the core (source-sink functioning). Finer population genetics and landscape analyses in North-East-France and in Belgium should enable to assess whether this pattern can be extrapolated to all colonies located on this northern limit of *R. ferrumequinum* distribution range or whether it is very special to this colony.

The last particular situation emphasised in this study concerned Tunisia. The level of genetic diversity detected in this colony was close to the levels observed in European mainland colonies, but private allele richness was higher. Moreover, this Tunisian colony exhibited a signature of demographic expansion, although we have to be cautious with this result as the confidence interval of the estimated time since the expansion started was extremely large (from 1 to 11,000 generations). Larger and denser sampling of colonies of *R. ferrumequinum* from North Africa would be necessary to assess whether it might suit to all colonies located on this southern part of the Mediterranean Sea.

### Implications for conservation

In this study we have shown that connectivity, genetic diversity levels and effective population size are high and homogeneous on the European mainland, when excluding the Northern French colony ‘MON’. Therefore the French Poitou-Charentes region does not need to be considered as a management Unit. We rather recommend considering the large population on the European mainland as a unique management unit. This large population could be resilient to local disturbance because of its strong interconnection between colonies. However we cannot exclude that some particular colonies within this population might be vulnerable, if exposed to human activities such as pesticides spraying or destruction of favourable habitats. It is therefore still important to pursue colony surveys at local scales, and also to standardize monitoring procedures at the national scale. Our sampling scheme did not enable us to identify the eastern boundaries of this population. It would require more sampling in Eastern France and in neighbouring countries (e.g. Belgium, Germany, Luxembourg, and Switzerland). In the future, this delineation and inferences of bat population demography might be of particular importance as *R. ferrumequinum* experienced severe declines there, so that some populations might be at higher risk of extinction and deserve special management attention (see Ransome and Hutson, 2000).

We have also shown in this study that peripheral colonies are genetically poorer than those at the core range, because of genetic drift, low gene flow and small effective population size. Thus, these colonies are more vulnerable to extinction and deserve particular management efforts. Interestingly, these colonies located at the edge of the species range are genetically divergent and may harbour some genetic and phenotypic variability that could be important for adaptation to global changes (Lesica & Allendorf, 1995). For example these colonies may play a key role in the face of climate change by facilitating species range shift (Rebelo, Tarroso, & Jones, 2010).

Lastly, our results advocate for paying particular attention to mating territories, and to movement pathways that enable extra-colony mating. Conservation programs should include the identification and protection of mating site and their associated habitats, and the maintenance of connected and structured semi-open habitats (i.e. mosaic landscapes of broadleaf woodland and grassland connected with tree lines) that are needed for bats to complete their yearly life cycle. Further collaborations between naturalist NGOs and academics are required to evaluate these relationships between landscape, movements at different scales, mating and genetic mixing, through the development of joint ecological and genetic approaches.

## Acknowledgments

First of all, we are indebted to all of our collaborators in the naturalist associations who helped us to generate this comprehensive bat sampling: Poitou-Charentes Nature (Deux-Sèvres Nature-Environnement, Nature-Environnement 17, la LPO France, Charente Nature et Vienne Nature), CREN Poitou-Charentes, Groupe Chiroptères des Pays-de-la-Loire, Groupe Chiroptères Aquitaine, Groupe Chiroptères du Groupe Mammalogique et Herpétologique du Limousin, Chauves-Souris Auvergne, Groupe Chiroptères de Provence, Groupe Mammalogique Normand, Amikiro, Coordination Mammalogique du Nord de la France, and all the numerous volunteers that contributed to fieldwork. Financial support was received from the LABEX ECOFECT (ANR-ll-LABX-0048) of Université de Lyon, within the program “Investissements d’Avenir” (ANR-ll-IDEX-0007) operated by the French National Research Agency (ANR). Orianne Tournayre PhD is funded by the LabEx CeMEB, an ANR “Investissements d’Avenir” program (ANR-10-LABX-04-01). Raphael Leblois was supported by the Agence Nationale de la Recherche (project GENOSPACE ANR-16-CE02-0008). Data were produced using the genotyping facilities on the GenSeq platform of LabEx CeMEB (Centre Méditerranéen Environnement et Biodiversitè). Part of this work was carried out by using the resources of the INRA MIGALE (http://migale.jouy.inra.fr) and GENOTOUL (Toulouse Midi-Pyrénées) bioinformatics HPC platforms, as well as the CBGP and the Montpellier Bioinformatics Biodiversity (MBB, supported by the LabEx CeMEB, an ANR “Investissements d’avenir” program ANR-10-LABX-04-01) HPC platform services.

## Data Archiving Statement

Data for this study are available at: to be completed after manuscript is accepted for publication.

## Supplementary material legends

Table SI. Microsatellites information. F= Forward, R=Reverse. Fam, Vic, Pet and Ned are the dyes used to make the PCR kits.

Table S2. Pairwise F_ST_ estimated overall loci between localities. Significant exact G-test after FDR correction between pairs of localities are represented with *** for pcO.OOl, ** for *p* <0.01 and * for *p* <0.05. NS indicates non-significant exact G-tests.

Table S3. Past demographic inferences using the OnePopVarSize model in MIGRAINE. ‘Pool_Englan? corresponds to the colonies from England and ‘Pool_France’ to all the French colonies. For each tested locality or pool of localities, the sample size (N), the value of the pGSM model parameter (pGSM), the estimated time at which the demographic change (i.e. bottleneck or expansion) happened (D), and the estimated intensity of the demographic change (Nratio) are represented. A contraction corresponds to a Nratio < 1 and an expansion to a Nratio > 1. The change in population size was determined significant, when 1 lies outside the 95% confidence intervals (95% CIs) of the Nratio. The demography event corresponds to the interpretation of these estimations. NS = Nonsignificant, S = significant, E = expansion, C = contraction.

Table S4. Estimation of the *ϑ* parameter which corresponds to 4*N_e_μ* using the OnePop model in MIGRAINE. ‘Pool_Englan? gather all the colonies from England and ‘Pool_continenta? all the French and Spanish Basque colonies. For each tested locality or pool of localities, the sample size (N), the value of the pGSM model parameter (pGSM), and the estimation the *ϑ* parameter. The population size was determined significant when 1 lies outside the 95% confidence intervals (95% CIs) of the *ϑ* estimates.

Figure S1. Determination of the most likelihood number of genetic clusters using the method proposed by Evanno et al. (2005). Likelihood of K, and ΔK are represented as a function of the number of groups K. ΔK estimates are based on the rate of change in the log probability of data between successive K values.

Figure S2. Evidence of genetic clusters was examined in a Principal Components Analysis(APC) using the R package ade4 (Dray and Dufour, 2007) and factoextra (Kassambara and Mundt, 2017). A) Percentage of explained variation for each principal component (left) and eigen values (right). B) Plot of the Principal Components Analysis.

Figure S3. Plot of the estimated θ by colonies in function of the distance to the centroid A) in France/Spanish Basque Country, B) in France/Spanish Basque Country/England. Red line represents the regression and its significance is indicated by the p-value above the graphs.

